# Pervasive chromatin remodeling at X-inactivation escape genes in schizophrenic males

**DOI:** 10.1101/300624

**Authors:** Hezha Hassan, Hedi Hegyi

## Abstract

Reanalyzing a large methylome dataset of 225 schizophrenic and 450 control samples derived from the prefrontal cortex revealed that 6 male patients have predominantly hypomethylated probes mostly on chromosome X, affecting the same genes in all six. Network analysis of the differentially methylated genes revealed a dense network of transcription factors, histone and chromatin remodeling proteins, with 15 of the X-located genes expressed at the synapse, including *NLGN4X*, *SYN1* and *MECP2.* Mapping a recent experimental dataset of G-quadruplexes (G4s) onto the differentially methylated probes revealed that the probes in the group of six overlapping with G4s on chromosome X are significantly more hypomethylated than non-overlapping and non-X probes whereas in the rest of the patients G4-overlapping probes are more methylated than non-overlapping ones, revealing a distinct pathology, involving chromatin remodeling for the six patients. Unexpectedly, the hypomethylated genes in them significantly overlapped with gene locations where X-inactivation escapism was observed in women.

## Introduction

Despite a large body of work and identification of more than 2000 genes implicated in schizophrenia, the exact cause of the disease has not been identified (1). Several genome-wide studies of differentially methylated genes in schizophrenics identified gene sets that are hypomethylated in some patient cohorts but appear to be normal or hypermethylated in others. Wockner et al. (2) using a commercial chip identified two subsets of patients, each consisting of 6 individuals, whose prefrontal cortex regions are differentially methylated in the opposite direction when compared to each other, comprising more than 10 thousand genes. Several other methylome studies have been carried out in schizophrenics. Pidsley et al. (3) found that the most enriched/differentially methylated genes in schizophrenics are associated with developmentally important genes, thus indirectly confirming the long-suspected notion that schizophrenia-related changes may start *in utero*.

Interestingly, despite a significant number of genes on chromosome X related to cognition and brain development (4, 5) there is only limited information that connects schizophrenia to X chromosome genes (6). In this study we reanalyzed a large set of methylomes acquired from the prefrontal cortex of 225 patient samples and 450 controls (7). We clustered the patient sample profiles using the numbers of differentially methylated probes for each gene and identified a cluster of six male patients that have their differentially methylated probes predominantly on chromosome X, the majority of which have been hypomethylated. Comparing the differentially methylated probes to that of the controls identified a smaller set of genes that may account for the disease in these patients.

Our analysis also revealed that the predominantly hypomethylated probes in the X chromosome cluster patients significantly overlap with genomic (DNA-based) G-quadruplexes, identified by a recent genome-wide experimental study by Balasubramanian et al (8). Comparing them to the rest of the patients in the Jaffe study (7) revealed that in those patients the genomic G-quadruplexes overlap mostly with hypermethylated probes, revealing a distinct pathology mechanism involving chromatin remodeling for the cluster of six patients.

Comparing the 68 genes with significant hypomethylation on chromosome X in the six patients with a list of 114 genes that tend to escape X-inactivation in women (9) revealed a significant overlap of 14 genes between the two sets, with even more genomic regions shared, raising the possibility that chromosome X-related chromatin remodeling contributes substantially to schizophrenia in a susceptible subset of patients.

## Results

In the first step we calculated the mean methylation values and their variance for each probe in the total of 287 adult control samples in the Jaffe study (7). We determined those probes in all the 225 patient samples that deviated from the control mean values by at least two standard deviations (Z-value >=2). We identified altogether 4883142 differentially methylated probes with an average number of 21703 differentially methylated probes per patient sample (all probe values deviating from the control means by at least 2 SDs are listed in **Supplementary Table 1**). Calculating the same for the controls resulted in 21822 probes per sample.

While the numbers are very similar for patients and controls, and the statistical significance for most of these probes does not reach genome-wide significance our goal at this step was the overall characterisation and clustering of patient profiles based on the similarities in methylation differences only, not the identification of driver genes with a clear-cut causative effect on schizophrenia.

### Chromosome-wide methylation values

After counting the differential methylation values for each sample and each chromosome (Supplementary table 2) we noticed that some of the patient samples (e.g. Sample250) have their highest number of hypomethylated probes on chromosome X. After comparing all patient sample profiles with one another we identified seven patient samples whose chromosome-wise counts of hypo- and hypermethylated probes correlated strongly within the group (Pearson correlation mean: 0.937) but much less between the group members and the rest of the samples (Pearson correlation mean: 0.578). One of the 7 samples, Sample596 was flagged by the original authors (7) as it failed a simple quality control, having been predicted male based on the sex chromosome probes, despite being derived from a female sample, therefore we ignored the sample in further analysis.

### Chromosome-wide methylation values in controls

Carrying out the same analysis of the differentially methylated probes in the controls (only the 287 adult control samples as the fetal samples showed vastly different methylation values as noted also by the authors in (7)) we found that two control samples (Sample273 and Sample506) also have the highest number of their hypomethylated probes on chromosome X. The chromosome-wise distribution of hypo- and hypermethylated probes in the two control samples on average correlated well with the cluster of 6 patients (r=0.89) but again much less with the rest of the patient samples (r=0.476).

### Exploring gene methylation profile similarities with Exploration Graph Analysis

To explore the methylation similarities gene-wise among the patients and controls we used a new method called EGA, Exploration Graph Analysis (10). EGA takes in an array of values for several samples and establishes the similarities among the samples by grouping them into a number of clusters.

Ranking both the patient and control samples for the hypomethylated probes on chromosome X, we selected the top 50 patient and top 30 control samples, containing both the cluster of six patient and the two control samples as outlined above. Using EGA for the hypomethylated probe counts of genes for these 80 samples resulted in seven groups (Fig 1). The green group consists of cluster 6 members and the two controls, clearly separated from the rest of the samples.

**Figure 1.**
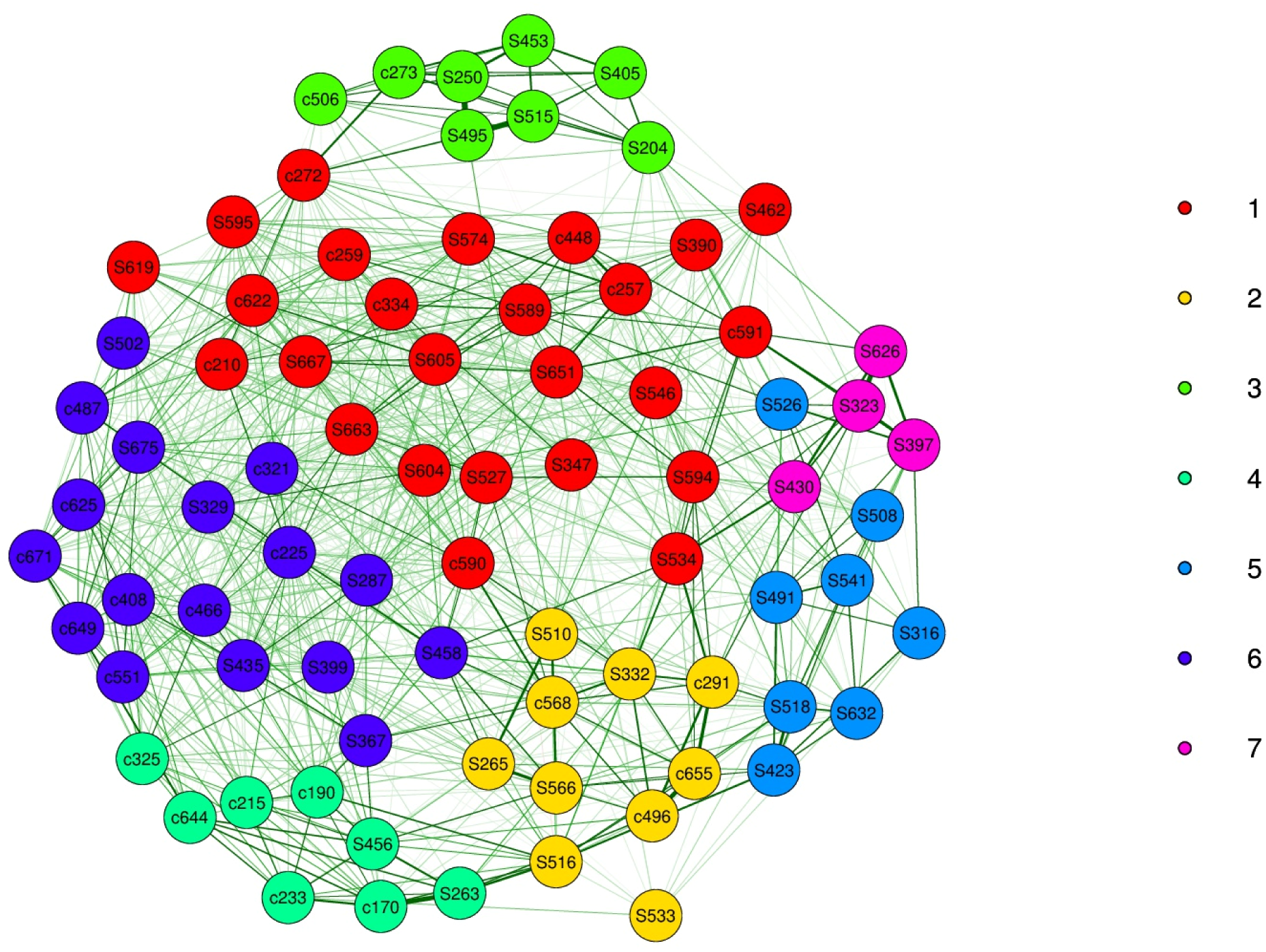
The 50 most hypomethylated patient samples and 30 most hypomethylated control samples clustered using EGA based on the counts of hypomethylated probes for all human genes in the Jaffe study (ref) deviating from control means by at least 3 SDs.

We also attempted to reconstruct the clusters of samples with EGA using only a reduced number of genes, using their counts of hypomethylated probes in the 50+30 (patient + control) samples. While using only chromosome X-coded genes did not recapture the same 6 patients +2 controls cluster, adding chromosome 1-coded genes (Supplementary Figure 1) resulted in the same grouping for the 6+2 cluster as the genome-wide probes in Figure 1.

### G-quadruplexes affect methylation differently in the cluster of six

In the next step we analyzed the differentially methylated probes in the cluster of six patients and two controls with relation to their overlap with G-quadruplexes. We used the G-quadruplex definitions established by Balasubramanian et al (8) downloaded from the Gene Expression Omnibus at https://www.ncbi.nlm.nih.gov/geo/query/acc.cgi?acc=GSE63874 and identified the probes on the Illumina chip that overlap with any of the G-quadruplexes in ref. (8) using the intersect function of bedtools. Figure 2 shows a boxplot statistics of the methylation of the CpG probes for G4-overlapping and G4-non-overlapping probes, separately for the six patients and the rest of them, and also for each group the chromosome X-located probes and the rest of the probes **(boxes A to H)**. Boxes **I to P** show the values for the controls, again separating the two controls (Sample273 and Sample506) with the hypomethylated X-chromosomes from the rest of the controls.

**Figure 2.**
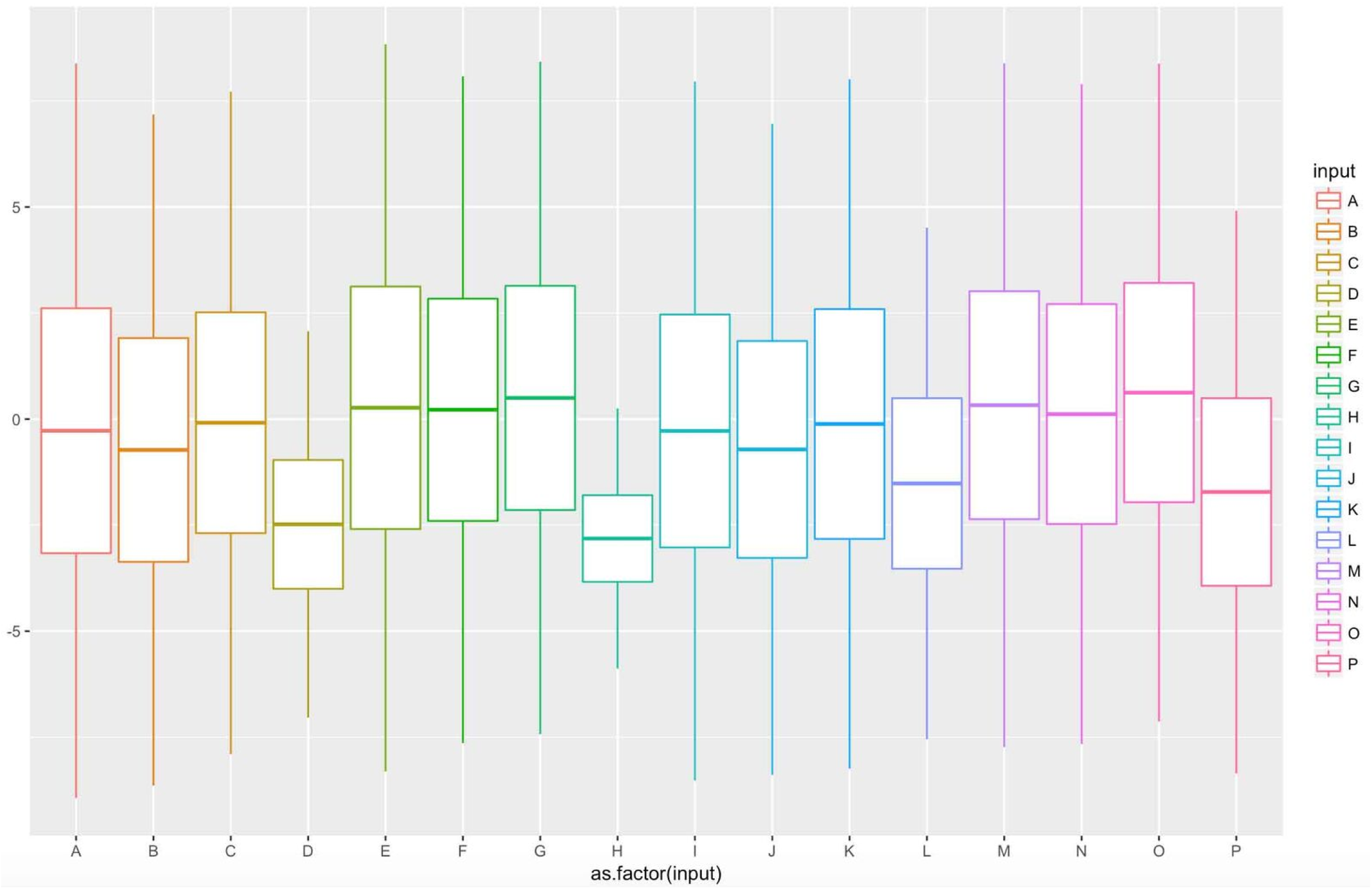
Boxplot statistics of mean methylation values for G4-overlapping (columns E to H and M to P) and non-overlapping CpG probes for patients (columns A to H) and controls (columns I to P). Patient columns: A & B: G4-non-overlapping, non-cluster 6, non-chromosome X and chromosome X probes, respectively; C & D: G4-non-overlapping, cluster 6, non-chrX and chrX probes, respectively; E & F: G4-overlapping, non-cluster 6, non-chrX and chrX probes, respectively; G & H: G4-overlapping, cluster 6, non-chrX and chrX probes, respectively. Columns I to P designations are the same as for A to H, but for the control samples. For the controls the cluster 6 is replaced with the two outlier controls, Samples 273 and 506.

We used Student’s t test to determine if any two sets of probes are statistically different. Chromosome X probes in the cluster 6 patients are apparently the most hypomethylated, the G4-overlapping probes being more hypomethylated (**box H**, mean: −2.815) than the non-G4-overlapping ones (**box D**, mean: −2.482, p-value=1.9e-12). This is in contrast with the rest of the patients where the G4-overlapping chromosome X-located probes (**box F,** mean: 0.219) are significantly more methylated than the non-overlapping probes (**box B**, mean: −0.728, p-value =0). The G4 non-overlapping probes in cluster 6 are also significantly more hypomethylated on chromosome X (**box D**, mean: −2.482) than the non-overlapping non-X probes (**box C**, mean: −0.087, p-value=0). The most striking difference is for the G4-overlapping probes in cluster 6 between chromosome X (**box H**, mean: −2.815) and non-X probes (**box G**, mean: 0.498, p-value=0). This is in stark contrast with the rest of the samples where there is no difference between the G4-overlapping X-probes (**box E**, mean: 0.266) and G4-overlapping non-X probes (**box F**, mean: 0.219). Finally, it is notable that for the cluster 6 patients the G4-overlapping non-X probes (**box G**, mean: 0.498) are significantly more methylated than the non-overlapping non-X probes (**box C**, mean: −0.087, p-value=2e-06), similarly to the non-cluster 6 patients (G4-overlapping, non-X: **box E**, mean: 0.266 vs. non-overlapping, non-X: **box A**, mean: −0.274, p-value=0.00037).

We repeated the calculations for the control samples, again handling the two controls with cluster 6-like behavior separately from the rest (values shown in **boxes I to P** in Figure 2). The overall values are similar in most cases, however there is a significant difference between cluster 6 of patients (columns 4, mean: −2.482) and the two outlier controls for the chromosome X probes (column 12, mean: −1.518) between the non-overlapping probes (p-val=0.002) but no significant difference for the G4-overlapping probes for chromosome X (columns 8 & 16, mean values: −2.815 & −1.719, p-value=0.14).

### Specific genes distinguishing the 6 patients from the controls

To pinpoint specific genes that distinguish the six patients from the controls we took into account only those probes that differ by at least 3 standard deviations (3 SDs) from the mean values derived from the controls.

After counting the differentially methylated probes using the 3 SD threshold for each gene and each sample we ran a t-test for each gene between the individual counts for the 6 patients and the controls (after removing the two outlier controls). We also calculated the mean person-wise counts for the cluster of six and the controls. Table 1 shows the results, listing the 82 genes with p-value<0.05 for the t-test and the relative person-wise probe counts >5, i.e. in cluster 6 the gene in question has at least 5 times more probes per person on average than in the controls.

**Table 1.**
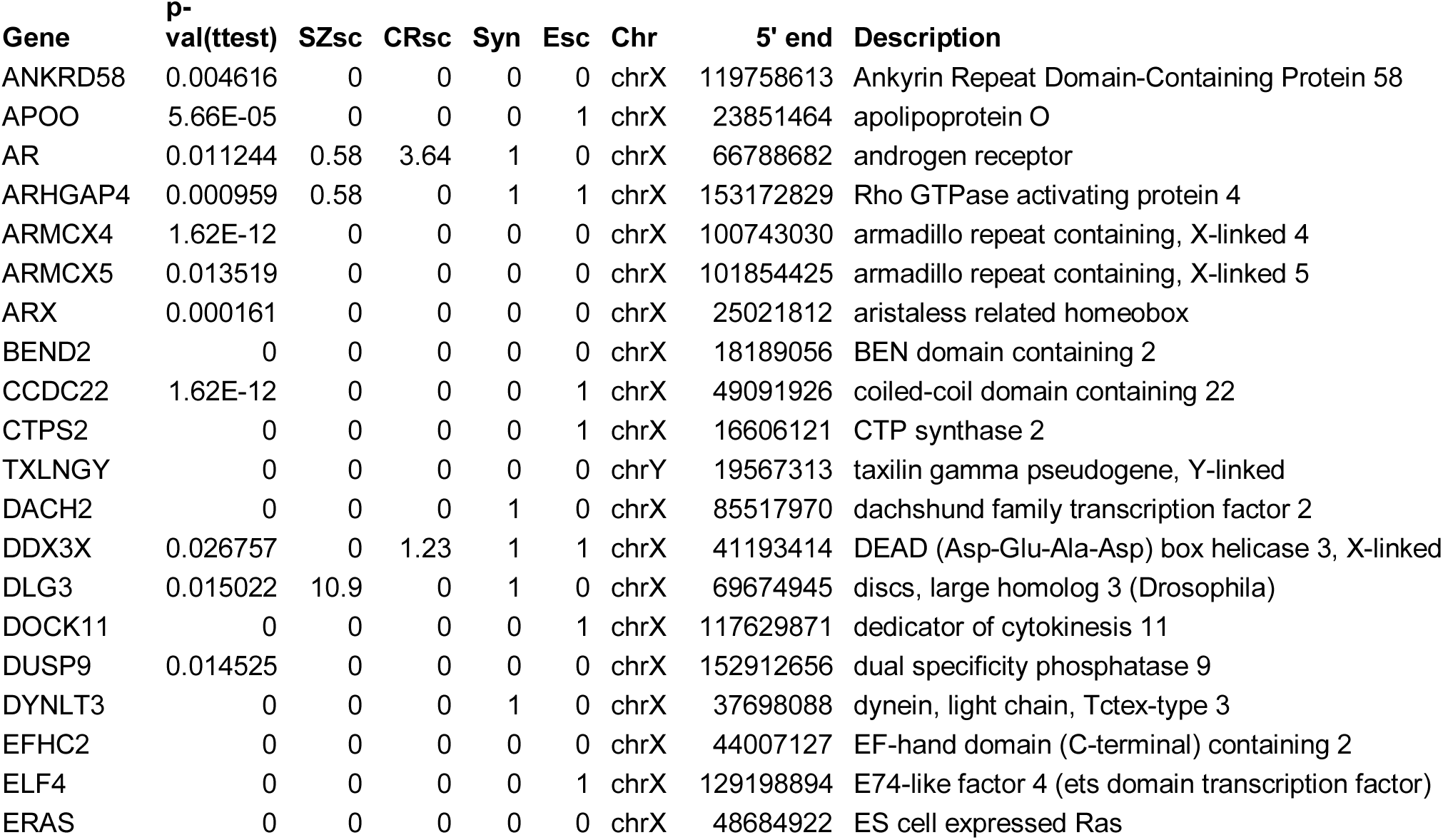
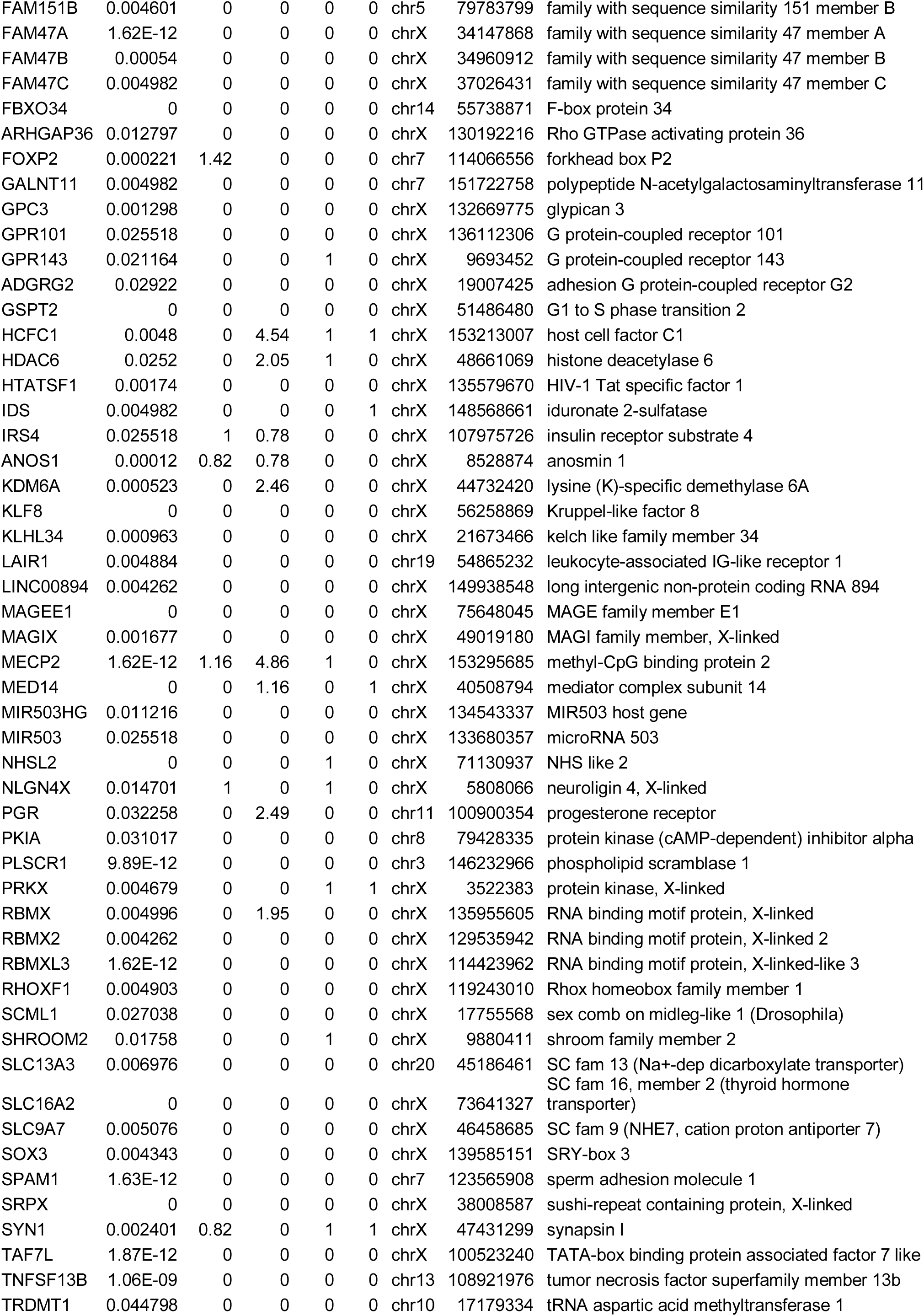
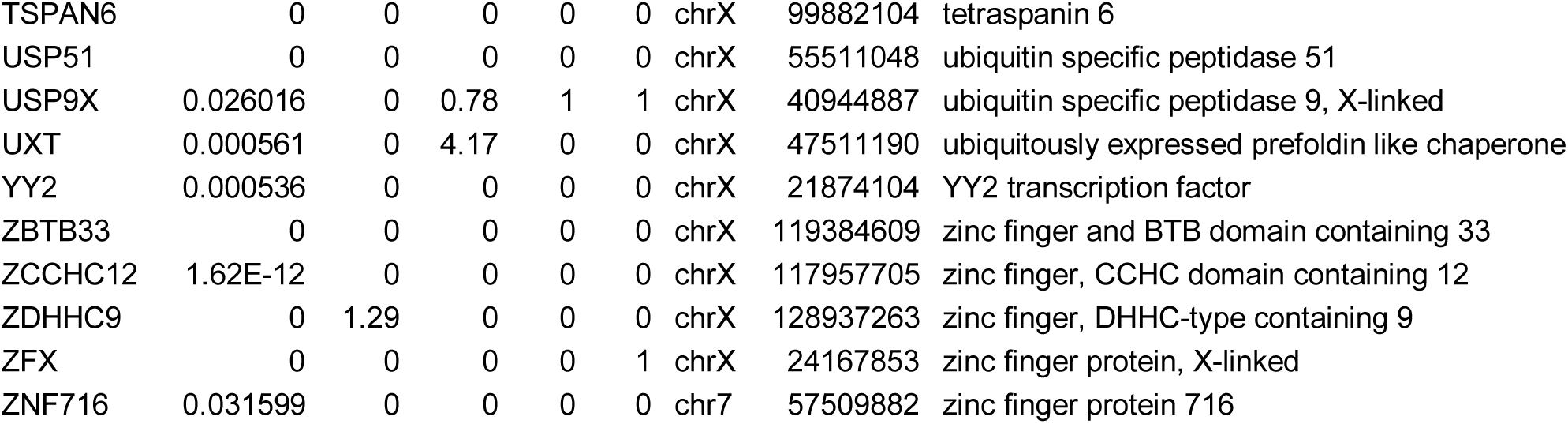
82 genes significantly hypomethylated (p-value<0.05) in the cluster 6 patients with relative person-wise counts of probes min. 5 times higher (in cluster 6) than in the control samples (except in the two outlier controls).

10 genes in the list (*RHOXF1*, *HTATSF1*, *ELF4*, *MED14*, *SOX3*, *TAF7L*, *DACH2*, *AR*, *ARX* and *YY2*) are transcription factors and altogether 27 genes (colored red in Fig 3) play a role in DNA-templated transcription. Three genes, *HDAC6*, *ZBTB33* and *MECP2* are related to histone deacetylation (4, 11, 12), *KDM6A* is a histone demethylase (13), while *UXT* is involved in chromosome remodeling (5). Two genes, *ZBTB33* and *MECP2* bind methylated CpG dinucleotides in chromosomal DNA (14, 15). In addition, *ZBTB33* promotes the formation of repressive chromatin structures (16). *MECP2* is also the key gene in Rett syndrome (17). Altogether these genes clearly indicate a dysfunctional transcriptional regulation, involving chromatin remodeling and abnormal methylation, the latter to be a well-known hallmark of schizophrenia.

**Figure 3.**
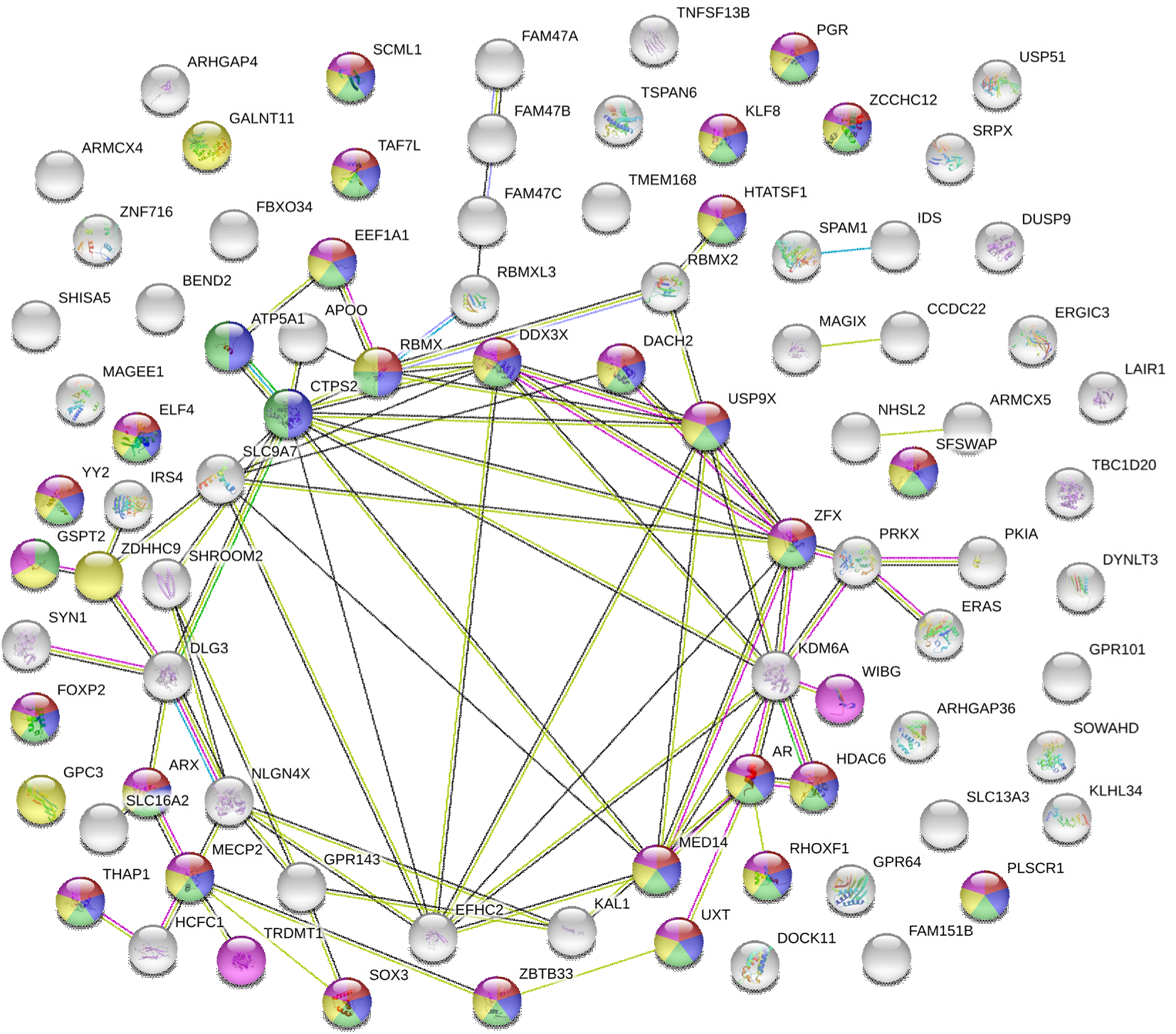
Protein-protein interactions (PPIs) among the protein-coding genes according to STRING (18). Proteins shown in red are involved in “DNA-templated transcription”, blue is for “macromolecule biosynthesis process”, green for “nucleobase-containing compound biosynthetic process and yellow is for “RNA metabolic process”. Most genes are located on chromosome X, 4 genes (*FOXP2*, *GALNT11*, *SPAM1* and *ZNF716*) on chromosome 7 and one pesudogene, *CYorf15A*, on the Y chromosome.

### Synaptic genes

The list also contains SYN1, synapsin I, a synaptic vesicle associated phosphoprotein, involved in the fine regulation of neurotransmitter release (19). According to Genecards (20), there are four more synaptic genes in our list: *DLG3*, *MECP2*, *NLGN4X* and *TSPYL2* (21–24). As schizophrenia is called the disease of the synapse (25), this finding warranted a more in-depth analysis. In a recent work Piton et al collected and resequenced all X-chromosome synaptic genes (26). After filtering our results with their list, we found that 15 genes, 1/5^th^ of those listed in Table 1 are synaptic genes.

Seven of the synaptic genes (*DLG3*, *MECP2*, *KAL1*, *SYN1*, *ARHGAP4*, *NLGN4X* and *AR*) are listed by Genecards as schizophrenia-related (27–33). *DLG3* is critical for synaptogenesis and required for learning most likely through its role in synaptic plasticity (34). Piton et al. found that *MECP2*, the main Rett syndrome gene, showed an accumulation of non-synonymous rare variants in schizophrenics (26). *KAL1* is the Kallman-syndrome gene (35), its relation to schizophrenia is so far largely hypothetical, based on a single patient with dual pathologies (29). *ARHGAP4*, the Rho GTPase Activating Protein 4 has been found to be associated with schizophrenia in the Han Chinese population (31). Interestingly the gene is in close proximity of *MECP2*, both located on chromosome Xq28 (31).

One of the synaptic genes with the most hypomethylated probes is *NLGN4X*, a neuroligin and synaptic cell-adhesion molecule that connects presynaptic and post-synaptic neurons at synapses (23). While it has more known association with autism (36) than with schizophrenia, a recent large-scale study by the Psychiatric Genomics Consortium identified the region as being one of 108 loci with highest numbers of mutations in schizophrenics (37). It is also a “hub” gene, having a very high number of, 3960, interacting partners in the brain (38), as is true of many of the synaptic genes (37, 38). *NLGN4X* has more than 20 times more interacting partners than the 2^nd^ highest ranking gene, *MECP2* in Table 1, with 178 interacting partners.

### Chromatin remodeling in the cluster of six

The most striking feature in the six patient samples is the remarkably similar profile of hypomethylated genes on chromosome X. Another characteristic is the apparently high occurrence of hypomethylated genes with functions leading to chromatin remodeling as 13 out of the 86 genes in our list of differentially methylated genes in the six patients are annotated with chromatin remodeling by Genecards (39–49). Many of the affected genes in Table1 show the hallmarks of chromatin regulation/remodeling, such as *MECP2*, *HDAC6*, a histone deacetylase and *KDM6A*, a histone demethylase. Sex hormone receptors (*AR* & *PGR*) are also known to influence chromatin remodeling by binding to chromatin (50, 51).

Several more genes in our list in Table 1 have chromatin remodeling functions, either directly recognized, such as for MED14, mediator complex subunit 14 (52), *UXT*, ubiquitously expressed prefoldin-like chaperone (40) or by association in proteomics studies, such as *IRS4*, insulin receptor substrate 4, *RBMX*, RNA binding motif protein, X-linked and *USP9X*, Ubiquitin Specific Peptidase 9, X-Linked (49). *EEF1A1*, elongation factor 1 alpha subunit and *ATP5A1*, a mitochondrial ATP synthase were linked to chromatin remodeling by an RNA interference screen (48).

Further evidence for chromatin remodeling on chromosome X of the 6 patients is provided by the high level of hypomethylation for probes overlapping experimentally verified G-quadruplexes on their X chromosomes whereas the opposite trend is observed for the rest of the patients and the rest of the chromosomes. As noted by others, G-quadruplexes may serve as genome regulators by playing a role in chromatin remodeling (53, 54).

### Genes overlapping with X-inactivation escape regions

We also compared our list of hypomethylated genes with a recent compendium of 114 genes for which the authors found experimental evidence that these genes tend to escape X-chromosome inactivation in women (9). Unexpectedly, 14 of the 68 chromosome X-located genes in our list in Table 1 are also listed in their compendium as X-escapee genes (Figure 4). What is more, most of the 68 genes are in the same broadly defined genomic regions outlined in (9). As noted by the authors, there is heterogeneity in escape between the studied European and Yoruban populations regarding the affected genes but there is a consensus for the genomic locations for the two groups. In Fig 4 where we showed the distributions of their 114 and our 66 genes, **respectively**, there is an overall consensus between the two groups of genes, with a Pearson correlation of 0.532 (p-value= 0.00041) for the gene occurrences in the 41 bins that we used to represent the layout of the two gene sets.

**Figure 4.**
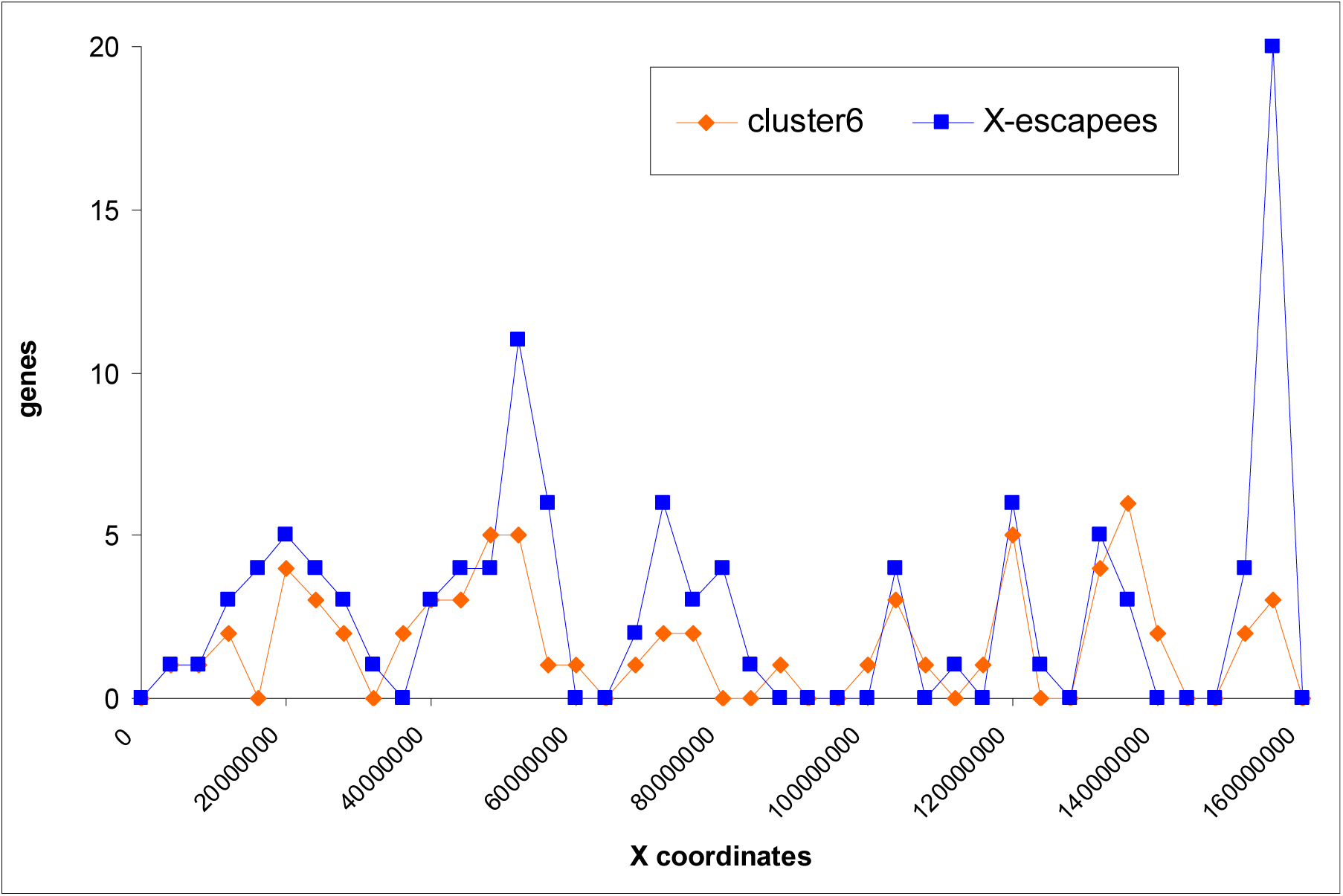
The distribution of the hypomethylated genes in the cluster of six male schizophrenics in (7) and X-escapee genes in (9) on the X chromosome in 41 equal-length (4 Mbase) bins.

### Comparing cluster 6 with two outlier controls

As two controls behave similarly to our cluster 6 samples, the question arises if we can distinguish the two groups based on the methylation values. We used another t-test when we compared the individual counts for each hypomethylated gene for the cluster 6 patients against the counts for the two controls. However, the result was significant only for 18 genes out of the 82 in Table1 (indicated in bold).

In the next step we compared the *hypermethylated* probes against both the two outlier controls and the rest of the controls, using again Student’s t-test. This gave a more consistent result (Table 2) as we found 9 genes (out of a total of 11 genes **in the two comparisons**) that were significantly more hypermethylated in cluster 6 in both comparisons. These genes have been incorporated already in Fig 3. Interestingly, two hypermethylated genes have protein-protein interactions with hypomethylated genes, such as *ATP5A1*, a mitochondrial ATP synthase also implicated in schizophrenia (55), which interacts with *CTPS2*, CTP synthase 2 and *WIBG*, a key regulator of the exon junction complex (56), which interacts with *KDM6A.* The antagonistic methylation differences for these two pairs raises the possibility that they have genetic, rather than protein-protein interactions.

**Table 2.**
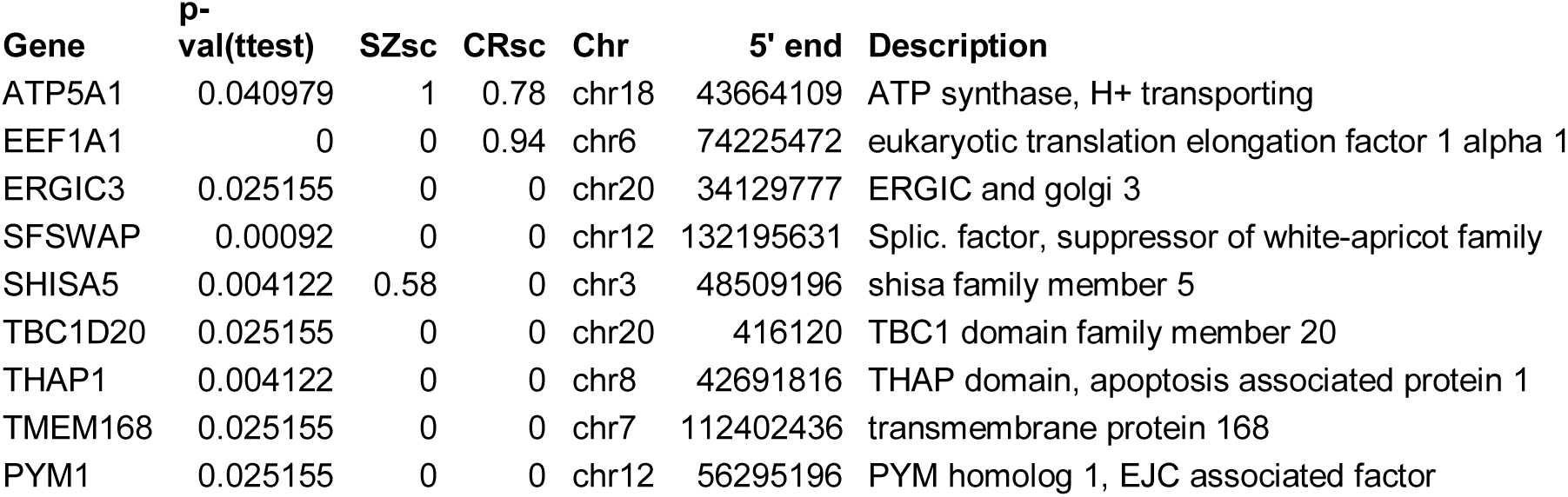
9 genes significantly hypermethylated (p-value<0.05) in the cluster 6 patients with relative person-wise counts of probes min. 5 times higher (in cluster 6) than in the control samples (except in the two outlier controls).

| Gene | p-val(ttest) | SZsc | CRsc | Chr | 5' end | Description |
| --- | --- | --- | --- | --- | --- | --- |
| ATP5A1 | 0.040979 | 1 | 0.78 | chr18 | 43664109 | ATP synthase, H+ transporting |
| EEF1A1 | 0 | 0 | 0.94 | chr6 | 74225472 | eukaryotic translation elongation factor 1 alpha 1 |
| ERGIC3 | 0.025155 | 0 | 0 | chr20 | 34129777 | ERGIC and golgi 3 |
| SFSWAP | 0.00092 | 0 | 0 | chr12 | 132195631 | Splic. factor, suppressor of white-apricot family |
| SHISA5 | 0.004122 | 0.58 | 0 | chr3 | 48509196 | shisa family member 5 |
| TBC1D20 | 0.025155 | 0 | 0 | chr20 | 416120 | TBC1 domain family member 20 |
| THAP1 | 0.004122 | 0 | 0 | chr8 | 42691816 | THAP domain, apoptosis associated protein 1 |
| TMEM168 | 0.025155 | 0 | 0 | chr7 | 112402436 | transmembrane protein 168 |
| PYM1 | 0.025155 | 0 | 0 | chr12 | 56295196 | PYM homolog 1, EJC associated factor |

We also observed an extremely hypermethylated single probe (cg00470565) in 4 of the 6 samples in our cluster, on chromosome 22 (Fig 5) deviating from the control mean by more than 9 SDs in all four of them. While the region is quite gene-rich, there is no gene annotation for this CpG probe. However, locating the probe in the UCSC genome browser (hg19, Fig 5) shows that it lies between an evolutionary highly conserved region and an open chromatin region with a high number of transcription factor binding sites.

**Figure 5.**
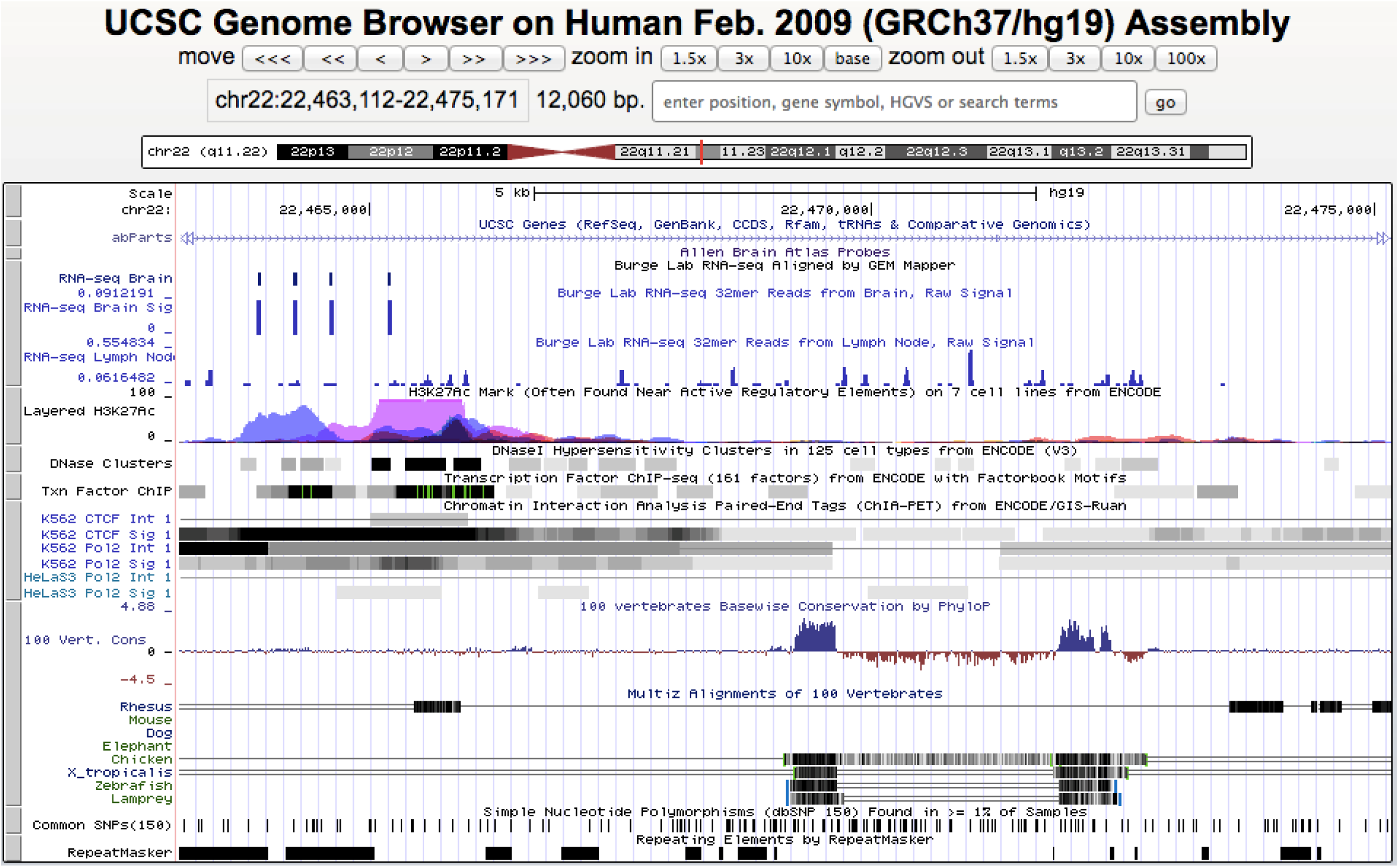
The genomic region around Illumina 450K probe cg00470565 on chromosome 22 at position 22,469,147 (using the coordinates of the human genome version hg19). There is no gene annotation for the probe but it lies in a highly conserved region, surrounded by open chromatin regions with a high number of transcription factor binding sites.

## Discussion

In this work we analyzed a recent large-scale methylome study of schizophrenics and controls where the authors queried the full methylome of 225 patient and 287 adult control samples taken from the prefrontal cortex, the brain region most implicated in schizophrenia.

We identified six male patient samples with a distinct methylation profile where the most differentially methylated probes were identified on chromosome X, most of which have been hypomethylated. We also identified two controls (one of them a female) with profiles very similar to the cluster of 6 but with less extreme hypomethylation values. While this chromosome X behavior may not necessarily lead to schizophrenia, it apparently predisposes the person to it, as the frequency ratio between schizophrenic and control population is more than 10:1 in males (restricting the calculations to adult males in the cohort in (7): 6 out of 119 patients vs. 1 out of 208 controls = 10.49).

Chromatin remodeling has been observed previously in neurodevelopmental and intellectual disorders (57), including schizophrenia (28), however it usually implicates one of four protein complex families as described in (58), none of which appears in our list of genes. However, the consistent hypomethylation of 68 X-coded genes in Table 1 in the six patient samples clearly indicates a pervasive chromatin remodeling of the X chromosome, apparently involving a new mechanism and new genes, different from the previously described ATPase-dependent chromatin remodelers (57).

The most unexpected result in our study is the apparent overlap (14 genes) between the hypomethylated genes in the six schizophrenic males and those that escape X-inactivation in women. As noted by Peeters et al. (59) X-inactivation has long served as a paradigm for heterochromatin formation, therefore the abnormal hypomethylation in the six schizophrenics is a further confirmation of chromatin remodeling in their X chromosomes. It shows that the same mechanism that lets the genes escape X-inactivation by unwinding the packaging of chromosome X in women makes it possible for these genes or genomic regions to escape epigenetic control in men, apparently leading or predisposing to schizophrenia.

X chromosome inactivation (XCI), a dosage compensation process in female mammals has been extensively studied in the last 50 years but still is not completely understood (60). The main factor is *XIST*, a long non-coding RNA that gradually coats and thus inactivates one of the two X chromosomes in females (59). It is known that about 15% of X-coded genes escape inactivation, which, however, vary both between and within species (9) but also share some common genes between human and mouse (59), some of which also appear in our list of hypomethylated genes (e.g. *DDX3X* and *KDM6A*). The exact cause of the escape from inactivation for these genes is not known but *XIST* has an antagonistic long noncoding RNA, *TSIX*, which raises the possibility that an escape from inactivation is also facilitated in part by long noncoding RNAs. We have the long noncoding RNA MGC16121/MIR503HG/H19X in our list, one of the most extensively hypomethylated genes in the six patients, which may have some role in the escape process. The gene was named H19X by Necsulea et al. (61) because it was co-expressed with H19, a conserved, maternally imprinted lncRNA (62) highly expressed in the brain. Several other lncRNAs have been suggested to play a role in X-inactivation and contribute to disease.

While we did not detect any aberrant methylation patterns in the six patients for the remodeling protein complexes mentioned above, we did find that X chromosome probes overlapping with G-quadruplexes (experimentally verified in (8)) are extremely hypomethylated in them whereas in other chromosomes in the same (as well as other) patients such probes tend to be more methylated than non-overlapping ones. This observation raises the possibility that chromatin remodeling in these patients is facilitated by these G-quadruplexes, acting perhaps as a molecular switch between euchromatin and heterochromatin. It has been recently observed that G-quadruplexes do play a role in the regulatory functions of chromatin (63).

As we also observed this phenomenon in two controls it raises the possibility that X chromosome remodeling may play a role in other diseases/syndrome especially when there is an observed difference between males and females such as autism, where the male: female occurrence ratio is about 3 (64). As we also observed the same X-related hypomethylation pattern in a female patient sample (but excluded it from further analysis as it did not pass quality threshold in the original publication), the possibility that the described X chromosome remodeling may cause schizophrenia also in women cannot be excluded.

## Methods

We downloaded methylation data accompanying ref. (7) from https://www.ncbi.nlm.nih.gov/geo/query/acc.cgi?acc=GSE74193. It contains methylation values of the Illumina 450K commercial chip querying 480 thousands locations across the human genome in the frontal cortex of 225 schizophrenic and 450 control samples. After removing samples with less than 17 years of age at the time of death we calculated statistics (mean, standard deviation (SD)) for the remaining 287 control samples for each of the 450k probes using an in-house Perl script (wherever not mentioned explicitly, we used such Perl scripts that are available from the authors).

### Clustering patients based on their hypo- and hypermethylated probe counts

We counted the differentially methylated probes for each chromosome for each patient and each adult control and compared the chromosome-wise profiles with one another calculating the Pearson correlation of the chromosome-wise counts, separately for the hypomethylated and hypermethylated probes (using the 2 SD thresholds to delineate the differentially methylated probes). We calculated Pearson correlations for every pair of samples of the 287 adult controls and the 225 patients using the chromosome-wise profiles of hypomethylated and hypermethylated probes using the 2SD thresholds.

We also counted the gene-wise differentially methylated probes for each sample. We took the 50 most hypomethylated patient samples (with the highest number of hypomethylated probes using the 2SD thresholds) and the 30 most hypomethylated control samples and generated clusters among them using Exploration Graph Analysis (EGA). EGA takes in an array of values for several samples and establishes the similarities among the samples by grouping them into a number of clusters. It is incorporated in R and Rstudio (10).

### Determining G-quadruplexes overlapping with hypomethylated probes

We used the G-quadruplex definitions established by Balasubramanian et al. (8) downloaded from https://www.ncbi.nlm.nih.gov/geo/query/acc.cgi?acc=GSE63874 and identified the probes on the Illumina 450K commercial chip that overlap with any of the G-quadruplexes in (8) using the intersect function of bedtools (39).

For Figure 2 we used the appropriate **Boxplot** R function.

## References

1. Mitchell KJ, Porteous DJ. Rethinking the genetic architecture of schizophrenia. Psychological medicine. 2011 Jan;41(1):19–32. PubMed PMID: 20380786.

2. Wockner LF, Noble EP, Lawford BR, Young RM, Morris CP, Whitehall VL, et al. Genome-wide DNA methylation analysis of human brain tissue from schizophrenia patients. Transl Psychiatry. 2014;4:e339. PubMed PMID: 24399042. Pubmed Central PMCID: PMC3905221.

3. Lam EY, Beraldi D, Tannahill D, Balasubramanian S. G-quadruplex structures are stable and detectable in human genomic DNA. Nature communications. 2013;4:1796. PubMed PMID: 23653208. Pubmed Central PMCID: 3736099.

4. Yoon HG, Chan DW, Reynolds AB, Qin J, Wong J. N-CoR mediates DNA methylation-dependent repression through a methyl CpG binding protein Kaiso. Molecular cell. 2003 Sep;12(3):723–34. PubMed PMID: 14527417.

5. Agger K, Cloos PA, Christensen J, Pasini D, Rose S, Rappsilber J, et al. UTX and JMJD3 are histone H3K27 demethylases involved in HOX gene regulation and development. Nature. 2007 Oct 11;449(7163):731–4. PubMed PMID: 17713478.

6. Ng MY, Levinson DF, Faraone SV, Suarez BK, DeLisi LE, Arinami T, et al. Meta-analysis of 32 genome-wide linkage studies of schizophrenia. Mol Psychiatry. 2009 Aug;14(8):774–85. PubMed PMID: 19349958. Pubmed Central PMCID: 2715392.

7. Jaffe AE, Gao Y, Deep-Soboslay A, Tao R, Hyde TM, Weinberger DR, et al. Mapping DNA methylation across development, genotype and schizophrenia in the human frontal cortex. Nat Neurosci. 2016 Jan;19(1):40–7. PubMed PMID: 26619358. Pubmed Central PMCID: PMC4783176.

8. Chambers VS, Marsico G, Boutell JM, Di Antonio M, Smith GP, Balasubramanian S. High-throughput sequencing of DNA G-quadruplex structures in the human genome. Nat Biotechnol. 2015 Aug;33(8):877–81. PubMed PMID: 26192317.

9. Zhang Y, Castillo-Morales A, Jiang M, Zhu Y, Hu L, Urrutia AO, et al. Genes that escape X-inactivation in humans have high intraspecific variability in expression, are associated with mental impairment but are not slow evolving. Molecular biology and evolution. 2013 Dec;30(12):2588–601. PubMed PMID: 24023392. Pubmed Central PMCID: 3840307.

10. Golino HF, Epskamp S. Exploratory graph analysis: A new approach for estimating the number of dimensions in psychological research. PloS one. 2017;12(6):e0174035. PubMed PMID: 28594839. Pubmed Central PMCID: 5465941.

11. Hubbert C, Guardiola A, Shao R, Kawaguchi Y, Ito A, Nixon A, et al. HDAC6 is a microtubule-associated deacetylase. Nature. 2002 May 23;417(6887):455–8. PubMed PMID: 12024216.

12. Theisen JW, Gucwa JS, Yusufzai T, Khuong MT, Kadonaga JT. Biochemical analysis of histone deacetylase-independent transcriptional repression by MeCP2. J Biol Chem. 2013 Mar 08;288(10):7096–104. PubMed PMID: 23349465. Pubmed Central PMCID: 3591619.

13. Lan F, Bayliss PE, Rinn JL, Whetstine JR, Wang JK, Chen S, et al. A histone H3 lysine 27 demethylase regulates animal posterior development. Nature. 2007 Oct 11;449(7163):689–94. PubMed PMID: 17851529.

14. Filion GJ, Zhenilo S, Salozhin S, Yamada D, Prokhortchouk E, Defossez PA. A family of human zinc finger proteins that bind methylated DNA and repress transcription. Mol Cell Biol. 2006 Jan;26(1):169–81. PubMed PMID: 16354688. Pubmed Central PMCID: 1317629.

15. Koch C, Stratling WH. DNA binding of methyl-CpG-binding protein MeCP2 in human MCF7 cells. Biochemistry. 2004 May 04;43(17):5011–21. PubMed PMID: 15109260.

16. Prokhortchouk A, Hendrich B, Jorgensen H, Ruzov A, Wilm M, Georgiev G, et al. The p120 catenin partner Kaiso is a DNA methylation-dependent transcriptional repressor. Genes & development. 2001 Jul 01;15(13):1613–8. PubMed PMID: 11445535. Pubmed Central PMCID: 312733.

17. Amir RE, Van den Veyver IB, Wan M, Tran CQ, Francke U, Zoghbi HY. Rett syndrome is caused by mutations in X-linked MECP2, encoding methyl-CpG-binding protein 2. Nature genetics. 1999 Oct;23(2):185–8. PubMed PMID: 10508514.

18. Szklarczyk D, Franceschini A, Wyder S, Forslund K, Heller D, Huerta-Cepas J, et al. STRING v10: protein-protein interaction networks, integrated over the tree of life. Nucleic acids research. 2015 Jan;43(Database issue):D447–52. PubMed PMID: 25352553. Pubmed Central PMCID: 4383874.

19. Greengard P, Valtorta F, Czernik AJ, Benfenati F. Synaptic vesicle phosphoproteins and regulation of synaptic function. Science. 1993 Feb 05;259(5096):780–5. PubMed PMID: 8430330.

20. Safran M, Dalah I, Alexander J, Rosen N, Iny Stein T, Shmoish M, et al. GeneCards Version 3: the human gene integrator. Database: the journal of biological databases and curation. 2010;2010:baq020. PubMed PMID: 20689021. Pubmed Central PMCID: 2938269.

21. Chen BS, Gray JA, Sanz-Clemente A, Wei Z, Thomas EV, Nicoll RA, et al. SAP102 mediates synaptic clearance of NMDA receptors. Cell reports. 2012 Nov 29;2(5):1120–8. PubMed PMID: 23103165. Pubmed Central PMCID: 3513525.

22. Na ES, Nelson ED, Kavalali ET, Monteggia LM. The impact of MeCP2 loss- or gain-of-function on synaptic plasticity. Neuropsychopharmacology: official publication of the American College of Neuropsychopharmacology. 2013 Jan;38(1):212–9. PubMed PMID: 22781840. Pubmed Central PMCID: 3521965.

23. Sudhof TC. Neuroligins and neurexins link synaptic function to cognitive disease. Nature. 2008 Oct 16;455(7215):903–11. PubMed PMID: 18923512. Pubmed Central PMCID: 2673233.

24. Wang GS, Hong CJ, Yen TY, Huang HY, Ou Y, Huang TN, et al. Transcriptional modification by a CASK-interacting nucleosome assembly protein. Neuron. 2004 Apr 08;42(1):113–28. PubMed PMID: 15066269.

25. Faludi G, Mirnics K. Synaptic changes in the brain of subjects with schizophrenia. International journal of developmental neuroscience: the official journal of the International Society for Developmental Neuroscience. 2011 May;29(3):305–9. PubMed PMID: 21382468. Pubmed Central PMCID: 3074034.

26. Piton A, Gauthier J, Hamdan FF, Lafreniere RG, Yang Y, Henrion E, et al. Systematic resequencing of X-chromosome synaptic genes in autism spectrum disorder and schizophrenia. Mol Psychiatry. 2011 Aug;16(8):867–80. PubMed PMID: 20479760. Pubmed Central PMCID: 3289139.

27. Clinton SM, Haroutunian V, Meador-Woodruff JH. Up-regulation of NMDA receptor subunit and post-synaptic density protein expression in the thalamus of elderly patients with schizophrenia. J Neurochem. 2006 Aug;98(4):1114–25. PubMed PMID: 16762023.

28. McCarthy SE, Gillis J, Kramer M, Lihm J, Yoon S, Berstein Y, et al. De novo mutations in schizophrenia implicate chromatin remodeling and support a genetic overlap with autism and intellectual disability. Mol Psychiatry. 2014 Jun;19(6):652–8. PubMed PMID: 24776741. Pubmed Central PMCID: 4031262.

29. Vagenakis GA, Hyphantis TN, Papageorgiou C, Protonatariou A, Sgourou A, Dimopoulos PA, et al. Kallmann’s syndrome and schizophrenia. International journal of psychiatry in medicine. 2004;34(4):379–90. PubMed PMID: 15825586.

30. Browning MD, Dudek EM, Rapier JL, Leonard S, Freedman R. Significant reductions in synapsin but not synaptophysin specific activity in the brains of some schizophrenics. Biol Psychiatry. 1993 Oct 15;34(8):529–35. PubMed PMID: 8274580.

31. Wong EH, So HC, Li M, Wang Q, Butler AW, Paul B, et al. Common variants on Xq28 conferring risk of schizophrenia in Han Chinese. Schizophr Bull. 2014 Jul;40(4):777–86. PubMed PMID: 24043878. Pubmed Central PMCID: 4059435.

32. Sand P, Langguth B, Hajak G, Perna M, Prikryl R, Kucerova H, et al. Screening for Neuroligin 4 (NLGN4) truncating and transmembrane domain mutations in schizophrenia. Schizophr Res. 2006 Feb 28;82(2-3):277–8. PubMed PMID: 16377159.

33. Culjkovic B, Stojkovic O, Savic D, Zamurovic N, Nesic M, Major T, et al. Comparison of the number of triplets in SCA1, MJD/SCA3, HD, SBMA, DRPLA, MD, FRAXA and FRDA genes in schizophrenic patients and a healthy population. American journal of medical genetics. 2000 Dec 04;96(6):884–7. PubMed PMID: 11121205.

34. Kumar R, Ha T, Pham D, Shaw M, Mangelsdorf M, Friend KL, et al. A non-coding variant in the 5’ UTR of DLG3 attenuates protein translation to cause non-syndromic intellectual disability. European journal of human genetics: EJHG. 2016 Nov;24(11):1612–6. PubMed PMID: 27222290. Pubmed Central PMCID: 5110046.

35. Franco B, Guioli S, Pragliola A, Incerti B, Bardoni B, Tonlorenzi R, et al. A gene deleted in Kallmann’s syndrome shares homology with neural cell adhesion and axonal path-finding molecules. Nature. 1991 Oct 10;353(6344):529–36. PubMed PMID: 1922361.

36. Landini M, Merelli I, Raggi ME, Galluccio N, Ciceri F, Bonfanti A, et al. Association Analysis of Noncoding Variants in Neuroligins 3 and 4X Genes with Autism Spectrum Disorder in an Italian Cohort. International journal of molecular sciences. 2016 Oct 22;17(10). PubMed PMID: 27782075. Pubmed Central PMCID: 5085789.

37. Schizophrenia Working Group of the Psychiatric Genomics C. Biological insights from 108 schizophrenia-associated genetic loci. Nature. 2014 Jul 24;511(7510):421–7. PubMed PMID: 25056061. Pubmed Central PMCID: PMC4112379.

38. Hegyi H. Connecting myelin-related and synaptic dysfunction in schizophrenia with SNP-rich gene expression hubs. Scientific reports. 2017 Apr 06;7:45494. PubMed PMID: 28382934. Pubmed Central PMCID: 5382542.

39. Quinlan AR, Hall IM. BEDTools: a flexible suite of utilities for comparing genomic features. Bioinformatics. 2010 Mar 15;26(6):841–2. PubMed PMID: 20110278. Pubmed Central PMCID: 2832824.

40. Liu L, McKeehan WL. Sequence analysis of LRPPRC and its SEC1 domain interaction partners suggests roles in cytoskeletal organization, vesicular trafficking, nucleocytosolic shuttling, and chromosome activity. Genomics. 2002 Jan;79(1):124–36. PubMed PMID: 11827465. Pubmed Central PMCID: 3241999.

41. Menon T, Yates JA, Bochar DA. Regulation of androgen-responsive transcription by the chromatin remodeling factor CHD8. Molecular endocrinology. 2010 Jun;24(6):1165–74. PubMed PMID: 20308527. Pubmed Central PMCID: 2875808.

42. Ceballos-Chavez M, Subtil-Rodriguez A, Giannopoulou EG, Soronellas D, Vazquez-Chavez E, Vicent GP, et al. The chromatin Remodeler CHD8 is required for activation of progesterone receptor-dependent enhancers. PLoS Genet. 2015 Apr;11(4):e1005174. PubMed PMID: 25894978. Pubmed Central PMCID: 4403880.

43. Mak AB, Ni Z, Hewel JA, Chen GI, Zhong G, Karamboulas K, et al. A lentiviral functional proteomics approach identifies chromatin remodeling complexes important for the induction of pluripotency. Molecular & cellular proteomics: MCP. 2010 May;9(5):811–23. PubMed PMID: 20305087. Pubmed Central PMCID: 2871416.

44. Li X, Wang W, Wang J, Malovannaya A, Xi Y, Li W, et al. Proteomic analyses reveal distinct chromatin-associated and soluble transcription factor complexes. Molecular systems biology. 2015 Jan 21;11(1):775. PubMed PMID: 25609649. Pubmed Central PMCID: 4332150.

45. Kanhoush R, Beenders B, Perrin C, Moreau J, Bellini M, Penrad-Mobayed M. Novel domains in the hnRNP G/RBMX protein with distinct roles in RNA binding and targeting nascent transcripts. Nucleus. 2010 Jan-Feb;1(1):109–22. PubMed PMID: 21327109. Pubmed Central PMCID: 3035123.

46. Rowbotham SP, Barki L, Neves-Costa A, Santos F, Dean W, Hawkes N, et al. Maintenance of silent chromatin through replication requires SWI/SNF-like chromatin remodeler SMARCAD1. Molecular cell. 2011 May 06;42(3):285–96. PubMed PMID: 21549307.

47. Lambert JP, Tucholska M, Go C, Knight JD, Gingras AC. Proximity biotinylation and affinity purification are complementary approaches for the interactome mapping of chromatin-associated protein complexes. J Proteomics. 2015 Apr 06;118:81–94. PubMed PMID: 25281560. Pubmed Central PMCID: 4383713.

48. Herr P, Lundin C, Evers B, Ebner D, Bauerschmidt C, Kingham G, et al. A genome-wide IR-induced RAD51 foci RNAi screen identifies CDC73 involved in chromatin remodeling for DNA repair. Cell discovery. 2015;1:15034. PubMed PMID: 27462432. Pubmed Central PMCID: 4860774.

49. Tsai YC, Greco TM, Boonmee A, Miteva Y, Cristea IM. Functional proteomics establishes the interaction of SIRT7 with chromatin remodeling complexes and expands its role in regulation of RNA polymerase I transcription. Molecular & cellular proteomics: MCP. 2012 May;11(5):60–76. PubMed PMID: 22586326. Pubmed Central PMCID: 3418843.

50. Urbanucci A, Marttila S, Janne OA, Visakorpi T. Androgen receptor overexpression alters binding dynamics of the receptor to chromatin and chromatin structure. The Prostate. 2012 Aug 01;72(11):1223–32. PubMed PMID: 22212979.

51. Vicent GP, Zaurin R, Nacht AS, Li A, Font-Mateu J, Le Dily F, et al. Two chromatin remodeling activities cooperate during activation of hormone responsive promoters. PLoS Genet. 2009 Jul;5(7):e1000567. PubMed PMID: 19609353. Pubmed Central PMCID: 2704372.

52. Wallberg AE, Yamamura S, Malik S, Spiegelman BM, Roeder RG. Coordination of p300-mediated chromatin remodeling and TRAP/mediator function through coactivator PGC-1alpha. Molecular cell. 2003 Nov;12(5):1137–49. PubMed PMID: 14636573.

53. Kejnovsky E, Lexa M. Quadruplex-forming DNA sequences spread by retrotransposons may serve as genome regulators. Mobile genetic elements. 2014 Jan 01;4(1):e28084. PubMed PMID: 24616836. Pubmed Central PMCID: 3933402.

54. Brazda V, Haronikova L, Liao JC, Fojta M. DNA and RNA quadruplex-binding proteins. International journal of molecular sciences. 2014 Sep 29;15(10):17493–517. PubMed PMID: 25268620. Pubmed Central PMCID: 4227175.

55. Altar CA, Jurata LW, Charles V, Lemire A, Liu P, Bukhman Y, et al. Deficient hippocampal neuron expression of proteasome, ubiquitin, and mitochondrial genes in multiple schizophrenia cohorts. Biol Psychiatry. 2005 Jul 15;58(2):85–96. PubMed PMID: 16038679.

56. Diem MD, Chan CC, Younis I, Dreyfuss G. PYM binds the cytoplasmic exon-junction complex and ribosomes to enhance translation of spliced mRNAs. Nature structural & molecular biology. 2007 Dec;14(12):1173–9. PubMed PMID: 18026120.

57. Lopez AJ, Wood MA. Role of nucleosome remodeling in neurodevelopmental and intellectual disability disorders. Frontiers in behavioral neuroscience. 2015;9:100. PubMed PMID: 25954173. Pubmed Central PMCID: 4407585.

58. Langst G, Manelyte L. Chromatin Remodelers: From Function to Dysfunction. Genes. 2015 Jun 12;6(2):299–324. PubMed PMID: 26075616. Pubmed Central PMCID: 4488666.

59. Peeters SB, Cotton AM, Brown CJ. Variable escape from X-chromosome inactivation: identifying factors that tip the scales towards expression. BioEssays: news and reviews in molecular, cellular and developmental biology. 2014 Aug;36(8):746–56. PubMed PMID: 24913292. Pubmed Central PMCID: 4143967.

60. Pinheiro I, Heard E. X chromosome inactivation: new players in the initiation of gene silencing. F1000Research. 2017;6. PubMed PMID: 28408975. Pubmed Central PMCID: 5373419.

61. Necsulea A, Soumillon M, Warnefors M, Liechti A, Daish T, Zeller U, et al. The evolution of lncRNA repertoires and expression patterns in tetrapods. Nature. 2014 Jan 30;505(7485):635–40. PubMed PMID: 24463510.

62. Brannan CI, Dees EC, Ingram RS, Tilghman SM. The product of the H19 gene may function as an RNA. Mol Cell Biol. 1990 Jan;10(1):28–36. PubMed PMID: 1688465. Pubmed Central PMCID: 360709.

63. Hansel-Hertsch R, Beraldi D, Lensing SV, Marsico G, Zyner K, Parry A, et al. G-quadruplex structures mark human regulatory chromatin. Nature genetics. 2016 Oct;48(10):1267–72. PubMed PMID: 27618450.

64. Loomes R, Hull L, Mandy WPL. What Is the Male-to-Female Ratio in Autism Spectrum Disorder? A Systematic Review and Meta-Analysis. Journal of the American Academy of Child and Adolescent Psychiatry. 2017 Jun;56(6):466–74. PubMed PMID: 28545751.

